# Neural Representation of the Relational Self from Infancy to Adulthood

**DOI:** 10.1101/2021.03.21.436295

**Authors:** Adi Ulmer-Yaniv, Shani Waidergoren, Ariel Shaked, Roy Salomon, Ruth Feldman

## Abstract

Investigations into the neural underpinnings of the “self” highlight its complexity and multi-dimensionality and emphasize that various aspects of the self are sustained by different neural systems. Here, we focused on the Relational Self, a dimension denoting the self-within-attachment-relationships that taps the continuity of attachment across individual development and affiliative bonds. Mothers and children were followed across two decades and videotaped in naturalistic interactions at three ages: infancy (3-6 months), childhood (9-12 years), and young adulthood (18-24 years). During fMRI scanning, young adults were exposed to videos of their own mother-child interactions from the three ages versus matched unfamiliar interactions. Relational Self-stimuli elicited greater activations across preregistered nodes of the human caregiving network, including thalamus-to-brainstem, amygdala, hippocampus, ACC, insula, and temporal cortex. Critically, Relational Self-stimuli were age-invariant in most regions of interest despite large variability of stimuli across multiple self-related features, such as similarity, temporal distance, affect, or mentalization, and Bayesian analysis indicated strong evidence for lack of age-related differences. PPI analysis demonstrated that Relational Self-stimuli elicited tighter connectivity between the ACC and insula. Greater child social engagement during interaction with mother correlated with higher ACC and insula response to Relational Self-stimuli. Findings highlight an important novel dimension in the neural representation of the self, suggest that the Relational Self may be sustained by a paralimbic interface integrating exteroceptive and interoceptive self-related signals, and demonstrate overlap in the attachment network of parents and children, lending support to perspectives on the continuity of attachment and self across the individual’s developmental history.

**Significance Statement:** Describing the neural underpinnings of the “self” is inherently complex due to the multi-dimensionality of the construct. Following mothers and children from infancy to adulthood, we focused on the Relational Self, a dimension denoting the self-within-attachment-relationships, and exposed young adults to own versus unfamiliar mother-child interactions across their relational history. Relational Self stimuli triggered greater activations in the human caregiving network, including thalamus-to-brainstem, amygdala, hippocampus, ACC, insula, and temporal cortex, were age-invariant, and elicited tighter connectivity between ACC and insula, creating a paralimbic interface of interoception-exteroception sustaining the Relational Self. Findings highlight a novel dimension in the neural representation of the self and lend support to perspectives emphasizing the cross-generational transmission of attachment and its continuity across the individual’s developmental history.

## Introduction

Representation of the “self” is an intriguing feature of human experience that combines the bodily (Blanke, 2012; Salomon, 2017), social (Decety and Sommerville, 2003; Yeshurun et al., 2021), and narrative (Christoff et al., 2011; Peer et al., 2015) aspects of the self and its ongoing transactions with the environment into a loosely-integrated construct (Northoff et al., 2006). While the neural underpinning of the self is a topic of continuous debate, current models postulate that its various dimensions are sustained by different cognitive processes and neural networks. For instance, the bodily self is thought to arise from the pre-reflexive integration of exteroceptive, interoceptive, sensory, and motor signals in fronto-parietal brain networks that sustain subjectivity in the present moment (Blanke et al., 2015; Park et al., 2016; Seghezzi et al., 2019). In contrast, episodic and narrative aspects of the self, which enable continuity of self-representation from the distant past to the projected future, rely on neural structures of the medial temporal lobe and default mode network (Andrews-Hanna et al., 2014). Still, across levels, the “self” is considered an organizational construct that maintains consistency across time and place; reflects personal valuation of goals and outcomes; attributes salience to internal and external cues; generates awareness of own body (Blanke et al., 2015); and guides interactions with conspecifics (Decety and Sommerville, 2003). It is also widely agreed that processes which underpin these fundamental models of the multi-dimensional “self” develop in infancy and early childhood in the context of the parent-infant attachment (Rochat, 2003; Feldman, 2017, 2020; Ciaunica and Crucianelli, 2019; Montirosso and McGlone, 2020). As such, the goal of the current study is to investigate the neural representation of the “Relational Self”, a dimension denoting the “self-within-attachment-relationships”, as measured across the individual’s attachment history with the mother from infancy to adulthood.

Studies on the brain basis of human attachment typically examined the parents’ neural response to their own infant stimuli as compared with unfamiliar infant, and recent studies exposed parents to naturalistic video vignettes of their own parent-child interaction compared with unfamiliar interaction (Noriuchi et al., 2008; Atzil et al., 2011; Abraham et al., 2014, 2016; Elmadih et al., 2016). Cumulative evidence from this line of research has delineated the neural structures that underpin human attachment (Feldman, 2015, 2017). The amygdala, ventral tegmental area (VTA), and hippocampus, areas rich in oxytocin receptors (Boccia et al., 2013; Raam et al., 2017), play a key role in mammalian mothering and are causally involved in bond formation (Insel and Harbaugh, 1989; Numan, 2020; Oxley and Fleming, 2000). In humans, these subcortical regions connect via multiple ascending and descending projections with several cortical regions to sustain the parent-infant attachment, particularly the anterior insula (AI), which supports interoception (Salomon et al., 2016) and empathy (Ulmer-Yaniv et al., 2020), the anterior cingulate cortex (ACC), a higher-order limbic interface for self-sustaining functions, and temporal regions of social observation and mentalization implicated in the reflective dimension of the self (Qin et al., 2020). Notably, fMRI studies of other human attachments throughout life, including romantic love and friendship, indicate that the same structures sustaining the parent-infant attachment also activate when individuals observe their romantic partner, co-parent, or close friends (Feldman, 2017; Bartels and Zeki, 2004; Acevedo et al., 2012; Abraham et al., 2017), suggesting continuity from parental to romantic to filial attachment. Yet, continuity in the neural response to cues representing the parent-child attachment from infancy to adulthood has not yet been examined.

Here, we utilized a longitudinal cohort followed from infancy to young adulthood. Naturalistic mother-child interactions were videotaped at three ages: infancy (3-6 months), childhood (10-12 years), and young adulthood (18-24 years). In young adulthood, participants were scanned in fMRI while observing these ecological vignettes spanning three stages in their Relational Self (i.e., Self) in addition to three vignettes of unfamiliar age-matched dyads (i.e., Other). Utilizing the same paradigm employed to study the neural basis of parental and romantic attachment, we compared neural activity during Self versus Other videos. We expected that the same structures that underpin parental and romantic attachment would also underlie the child’s neural response to attachment-related stimuli, including the amygdala, hippocampus, VTA, temporal cortex, insula, and ACC (Feldman, 2017) and that these regions would show higher BOLD activity when viewing Relational Self-stimuli compared with similar videos of unfamiliar dyads. In line with the formulations of attachment theory (Bowlby, 1969), we examined whether these regions would show a consistent, age-invariant responses to Relational Self-stimuli from different stages across the individual’s attachment history spanning two decades. Additionally, in light of research on the brain basis of the parent-child attachment (Leibenluft et al., 2004), we explored whether stimuli depicting the Relational Self would elicit increased functional connectivity among nodes of the attachment network. Finally, we tested whether the strength of activation in response to Self-stimuli would correlate with the degree of maternal sensitivity and child engagement in the interaction, two key behavioral markers of the mother-child attachment that have shown individual stability across development and support the child’s resilience, well-being, and adaptation (Feldman, 2010; 2021; Ulmer-Yaniv et al., 2018).

## Materials and Methods

### Participants

Participants included 65 young adults (mean age=20.03 years, SD=2.0, 33 males) who were followed from infancy and participated in two visits at the adult stage, a home visit and a brain imaging session. All participants were healthy, without chronic medical or psychiatric conditions, and completed at least 12 years of education. All children were born to middle-class, two-parent families, and their parents completed high-school education, were above 21 years at the child’s birth, were above poverty line, and were screened for psychiatric or psychosocial conditions. All participants were of Israeli-Jewish ethnicity.

### Longitudinal study design

Families were recruited in infancy to participate in a longitudinal study on mother-infant attachment and its developmental consequences. Home visits were conducted at three time-points; Infancy (3-6 months), childhood (9-12 years), and young adulthood (17-25 years). Visits were scheduled for the afternoon or evening hours. We were specifically interested in observing naturalistic interactions between mother and child which reflect as much as possible their habitual, daily interaction. For all interactions, mother and child were videotaped in a face-to-face position. The cameras were placed at 1.2 meter from the interacting dyad and filming tried to capture participants’ faces as much as possible.

At the adult stage, the two visits were conducted within a ∼6-week period of each other; Home visits were conducted in the evening hours and lasted ∼3.5 hours and were followed by MRI scan session. The study was approved by the Bar-Ilan University’s IRB. Both mother and young adult signed informed consent and received a gift card of 250 NIS (∼60$) for their participation.

Our pre-registered study included two cohorts. Cohort 1 was used for ROI definition and included 15 participants who were randomly assigned to this cohort 1. Cohort 2, our main study, included 50 participants with a full data set. Results reported here are based on data from these fifty participants.

81 participants had data from all three stages of the study, including the MRI scan. 10 subjects were discarded due to misplacement in scanner,1 for no visual activity and one for bad T1 scan. 4 additional subjects were discarded due to problems with the video stream during scan (video stuck, or mistakenly doubled). No differences on demographic information emerged between participants with and without valid data.

### Home observation of mother-child interactions

In infancy (mean age = 4.8 months, SD = 1.1), mother and infant were videotaped at home and instructions were “play with your infant as you normally do” for five minutes. In late Childhood (mean age = 10.9 years, SD = 1.2) and young adulthood (mean age =20.03 years, SD = 2), mother and child engaged in a validated conversation-based positive interaction for seven minutes (Feldman et al., 2014; Ulmer-Yaniv et al., 2017, 2018). Interactions were used for offline coding and as fMRI stimuli. For the fMRI task, two minutes from the middle of the interaction were selected.

### Coding of social behavior

Videos of mother-child interactions were coded with the Coding Interactive Behavior (CIB) manual (Feldman, 1998). The CIB is a global rating system for social interactions with multiple codes for parent, child, and dyadic behavior and several manuals for coding social behavior from newborns to adults based on the same codes, whenever possible, and similar conceptual principles. The CIB was validated in hundreds of studies with infants, children, adolescents and adults across multiple cultures and high-risk conditions. The system has good psychometric properties, including construct validity, test-retest reliability, and predictive validity (Feldman, 2012, 2021).

### In the current study we used two constructs

*Maternal sensitivity* – the averaged codes of mother’s acknowledgement of child communication, constant gaze, warm positive affect, warm vocalization, appropriate range of affective expression, and consistent style from the three ages (alpha = .92).

*Child Positive Engagement*, which averaged codes related to social engagement, affection and trust toward parent, positive affect, and involvement from the three stages (alpha = .86). The construct includes only child, not adult behaviors. In infancy, codes included child positive affect and child social alertness. In late childhood and young adulthood, in addition to these codes, the construct also includes the following codes: child affection to parent, child trust and openness to parent, child involvement in the conversation, and child warmth.

Coding was conducted by two coders blind to other information and trained to 90% reliability. Inter-rater reliability, computed for 20% of the videos averaged 94% (k = .87).

### MRI data acquisition

Magnetic Resonance Imaging (MRI) data was collected using a 3T scanner (SIEMENS MAGNETOM Prisma syngo MR D13D, Erlangen, Germany) located at the Tel Aviv Sourasky Medical Center. Scanning was conducted with a 20-channel head coil for parallel imaging. Head motion was minimized by padding the head with cushions, and participants were asked to lie still during the scan. High resolution anatomical T1 images were acquired using magnetization prepared rapid gradient echo (MPRAGE) sequence: TR=1860ms, TE=2.74ms, FoV=256mm, Voxel size=1×1×1mm, flip angle=8 degrees. Following, functional images were acquired using EPI gradient echo sequence. TR=2500ms, TE=30ms, 42 slices, slice thickness=3.5mm, FOV=220mm, Voxel size=3.2×2.3×3.5mm, flip angle=82 degrees. In total 393 volumes were acquired over the course of the empathy paradigm. Visual stimuli were displayed to subjects inside the scanner, using a projector (Epson PowerLite 74C, resolution = 1024 × 768), and were back-projected onto a screen mounted above subjects’ heads, and seen by the subjects via an angled mirror. The stimuli were delivered using windows media player software (Microsoft corporation). Before participating, participants signed an informed consent according to protocols approved by ethics committee of the Tel-Aviv Sourasky Medical Center. The study was approved by the Bar-Ilan University’s IRB and by the Helsinki committee of the Sourasky medical center, Tel Aviv (Ethical approval no. 0161-14-TLV). Subjects received a gift certificate of 300 NIS (∼85 USD) for their participation in the scan session.

### Self in relationship paradigm

In the Relational Self fMRI paradigm subjects were presented with a series of the video vignettes while lying in the scanner. “Self” Stimuli included a 2-minute movie of the subject-mother dyad interaction from each age period: infancy, childhood and young adulthood. In the “Other” condition participants viewed an unfamiliar mother and child dyad, matched for gender and age properties in a similar interaction. Stimuli were tailor-made for each subject in the home setting and the videos thus differed in various color, luminance, and sound properties. The order of presentation was counterbalanced so that half the subjects viewed themselves first, and half watched the strangers’ dyad first. (see figure 1 for details); however, the order of age conditions presentation was not randomized in order to present a coherent narrative on the maturation of the Relational Self from infancy to adulthood. The paradigm was developed in our lab and several studies with other populations were previously published (Atzil et al., 2011; Abraham et al., 2014)

**Figure 1:**
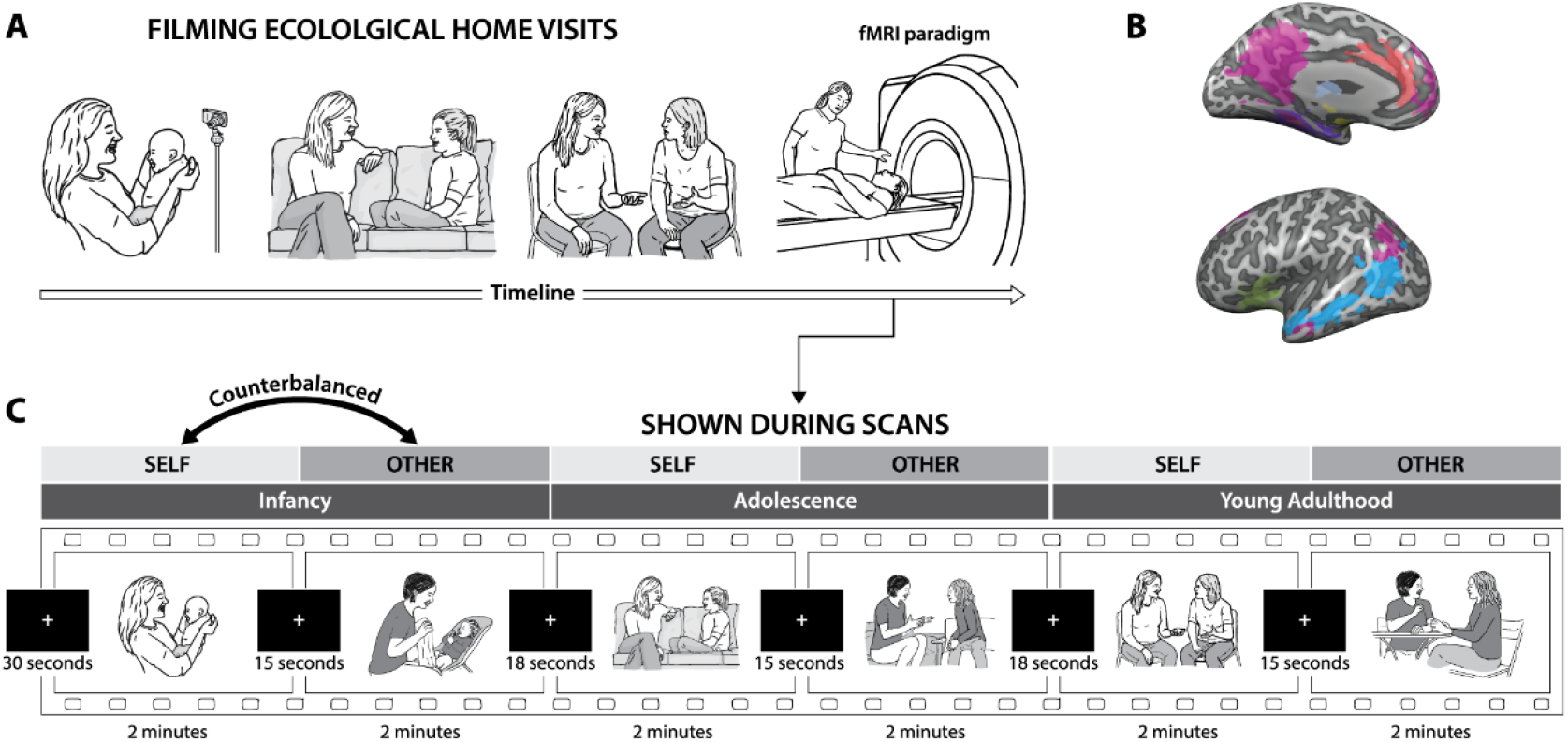
Longitudinal Research plan and fMRI paradigm. **A. Experimental procedure**. Participants were initially recruited as infants, around 3 months of age, participated again as children and later in young adulthood (current study phase). All sessions included a home visit in which a videotaped interaction of participants and their mothers took place. Video vignettes of interactions were used as fMRI stimuli. **B. Pre-registered regions of interest**: ACC (red), Insula (green), PHG (purple), Amygdala (yellow), temporal cortex (light blue) and DMN (fuchsia). **C. Experimental paradigm**. Participants were presented with six video vignettes of self and other dyadic interactions. Each age (infancy, childhood, young adulthood) was presented as an interaction of the participant and his mother (or a gender-matched stranger). Clips lasted 2 minutes each and were previewed by a fixation cross for 30 seconds. Between videos, fixation cross was presented for periods of alternately 15-18 seconds. Order of self-other was counterbalanced between participants.

### Data Analysis

### Data preprocessing

Data preprocessing and data analysis were conducted using BrainVoyager QX software package 20.6 (Brain Innovation, Maastricht, The Netherlands). The first 3 functional volumes, before signal stabilization, were automatically discarded by the scanner to allow for T1 equilibrium (resulting in 277 volumes). Preprocessing of functional scans included 3D motion correction, slice scan time correction and spatial smoothing by a full width at half maximum (FWHM) 6-mm Gaussian kernel. The functional images were then superimposed on 2D anatomical images and incorporated into the 3D datasets trilinear interpolation. The complete dataset was normalized into MNI space, using ICBM-452 template.

### Whole Brain Analysis

Multi-subject general linear model (GLM) was computed with random effects, with separate subject predictors, in which the different conditions (videos or fixation) were defined as predictors and convoluted with a standard hemodynamic response predictor. Following, a whole brain, two factors (*Self-relatedness (Self/Other)* \**Age(Infancy/ Childhood/Young Adulthood)*) repeated measures ANOVA was performed. Whole brain maps were created and corrected for false discovery rate (FDR) of q<0.050 (Benjamini and Hochberg, 1995). For visualization of results, the group contrasts were overlaid on a MNI transformed anatomical brain scan of a single participant.

### Regions-of-Interest Definition and Preregistration

Based on cohort 1 and a-priori theory-based selection, eight regions of interest were selected and pre-registered at Open Science Framework: Hippocampus and Parahippocampal gyrus (PHG), Amygdala, VTA, Anterior Cingulate cortex (ACC), Insula, a temporal cluster encompassing the superior temporal sulcus and gyrus, from the occipito-parietal border to the temporal pole, and the default mode network (DMN) as a network. Pre-registration is available at: https://osf.io/2ndxr/?view_only=ba738b07cad249e0b1f08c2f458ddb35

Cohort 1 included a group of 15 subjects (mean age 18.93 years old (SD=0.88), 46.7% males, 86.7% right-handed). Fixed effects multi subject GLM activation maps were used for ROI definition of the amygdala and thalamus. The DMN network was defined based on individual functional connectivity maps with seed in the precuneus, which were superimposed to create a 70% mutual probability map. The temporal cortex region was defined based on the pilot map combined with STS region from the Glasser atlas (Glasser et al., 2016). The insula, ACC, hippocampus-PHG were also taken from the Glasser atlas. VTA was defined by three 5mm spheres based on coordinates from the literature (Murty et al., 2014). Figure 2 shows the ROIs and figure S1 cohort 1 self>other map.

**Figure 2:**
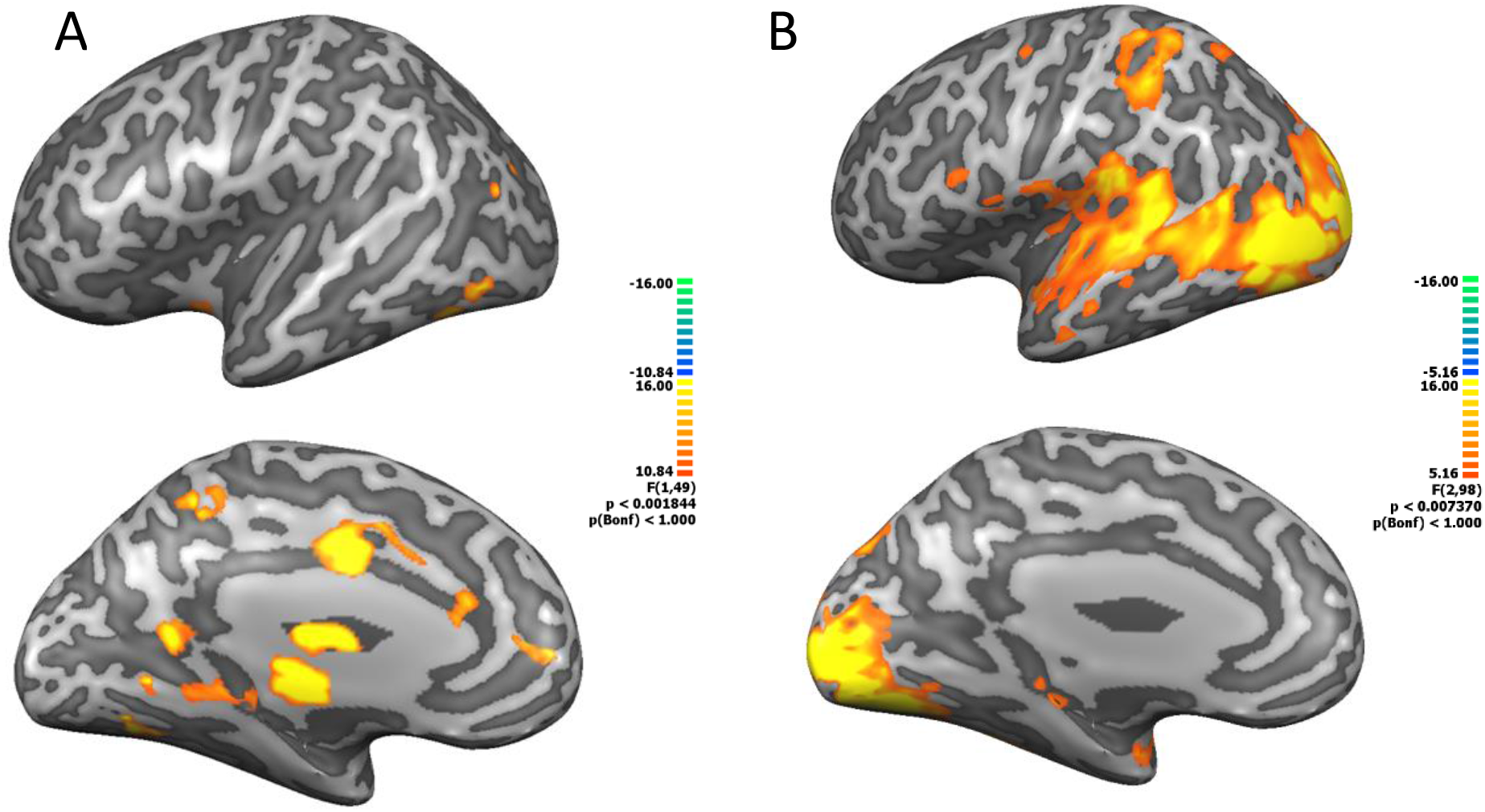
Whole brain two factorial repeated measures ANOVA. (Self/other*time (infancy, childhood, young adulthood). **A**: Main effect for Self/other, clusters in the ACC, thalamus and midbrain **B**: Main effect for time. Clusters in occipital regions, temporal lobe, parietal cortex and limbic regions. There was no Self/Other-Time interaction effect. All maps were FDR corrected(q(FDR)<0.05) on VMR and projected into an inflated brain for presentation purposes

### Psycho-physiological interaction (PPI) Analysis

Classic PPI analysis (Friston et al., 1997) was done using PPI plugin for BrainVoyager (V1.30) to PPI predictors and confounds as following: pre-registered ACC was defined as the seed region, and psychological conditions were defined as self>other, for all timepoints. For each condition, weight was assigned in such a way that the resulting time course will be zero centered (self adult+1, self child +1, self infant +1, other adult -1, other child -1, other infant -1). Fixation weight was set to zero.

For each subject, the time course of the ACC ROI was extracted, Z-transformed and then convolved with the hemodynamic response function. Following, it was multiplied TR by TR with the task time course (task time course was based on the protocol associated with the data) to create the ACC PPI predictor. Additionally, for each subject, a psychological regressor was created, based on the associated protocol, an ACC predictor, based on the ACC time course correlation and a complementary regressor. Additional motion correction predictors were added and Z-transformed. The resulting set of 4 PPI predictors for each subject, were used in a multi subject GLM analysis. The PPI ACC predictor allows to create a group map of voxels that increase their interaction with the ACC for the self conditions compared to other conditions, over and above what is explained by the task itself (*Self*>other contrast; psychological component) and by the global functional connectivity of the ACC (physiological component). Multi subject GLM analysis was restricted to our pre-registered ROIs, using a mask. Following, the ACC-PPI maps were corrected using Monte-Carlo cluster level statistical threshold estimator, with 1000 simulations to estimate cluster level probabilities (Forman et al., 1995).

### Statistical Analysis

Statistical analysis was conducted using JASP (Version 0.12.1 for windows, JASP Team, 2020), SPSS (SPSS statistics V25, IBM Corp.) and in R version 4.0.0 (R Core Team, 2020) with Tidyverse package (Wickham et al., 2019). Null effects were assessed using Bayesian statistics (Keysers et al., 2020). Greenhouse-Geiser correction was used for sphericity violations. Repeated measures Bayesian ANOVA was used to evaluate the evidence for the null effect found using the standard RM ANOVA analysis. We used the exclusion Bayes factor (BF_exc_). As such, a low value for BF_exc_ signifies support for the inclusion of the effect (i.e. evidence for the effect): BF_exc_ factors < 0.33 denote moderate evidence, BF_exc_ <0.1 denote strong evidence and BF_exc_ <0.03 denote very strong evidence for the inclusion of the model (Kelter, 2020).

## Results

As a first step, we examined the overall brain response to the video stimuli of the naturalistic mother-child interactions vs. the baseline fixation condition. A whole brain map, comparing the epochs of audio-visual stimulation to rest of cohort 1 dataset (15 subjects; figure S1A) and a similar map of the analysis dataset (50 subjects; figure S1B) are parallel. As expected, both maps show wide activations in the visual cortex, spreading to the temporal cortex. Additional activations were observed in the DMN, and in limbic regions; PHG, and amygdala. Note that the two maps were highly similar despite differences in the number of subjects.

Next, we examined the experimental factors using two factorial ANOVA analysis (Self-relatedness (*Self/Other*) *Age (*Infancy/Childhood/Young adulthood*) on whole brain activity (N=50). The ANOVA map of the *Self-relatedness* main effect (figure 2A) reveals that the main regions showing differential BOLD responses between the Self and Other conditions include the ACC, thalamus, and midbrain. Investigation of *Self>Other* contrast map indicated stronger activity for the *Self* condition in the middle ACC (BA 32), posterior ACC (BA 24), a large cluster spanning from the thalamus ventrally to the brain stem, and bilateral activations in the visual association regions – peristriate cortex (BA 19), SMA (BA 6) and cerebellum. In addition, a bilateral deactivation was found in the hypothalamus (figure S2, table 1).

**Table 1:**
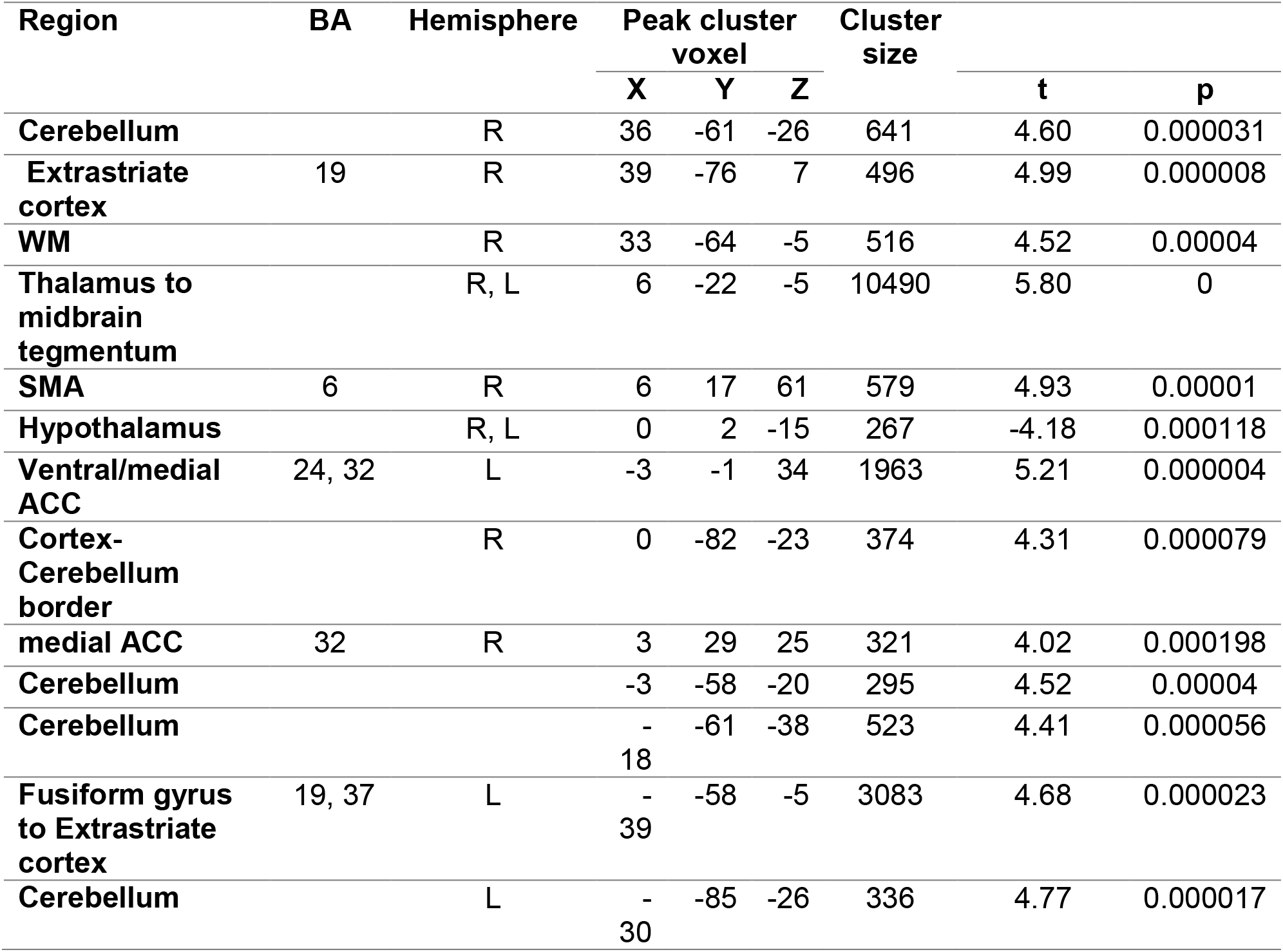
Clusters activated in Self>other contrast, analysis dataset.

**Table 2:** Clusters of PPI with ACC seed self>other map, restricted to pre-registered ROIs:

|  | Correlation | Preregistered ROI | BA | Hemisphere | Peak cluster voxel |  |  | cluster size | t | p |
| --- | --- | --- | --- | --- | --- | --- | --- | --- | --- | --- |
|  |  |  |  |  | X | Y | Z |  |  |  |
| Positive |  | Insula | 13 | L | -27 | 8 | -14 | 550 | 4.51 | 0.00004 |
|  |  | Temporal cortex | 22 | L | -48 | -7 | -17 | 110 | 3.43 | 0.001252 |
|  |  | PHG |  | L | -37 | -25 | -17 | 81 | 3.04 | 0.003801 |
|  |  | Amygdala | 53 | R | 33 | -1 | -20 | 42 | 2.85 | 0.006412 |
|  |  | VTA |  | L | -2 | -13 | -11 | 28 | 3.39 | 0.001379 |
|  |  | Insula | 47 | R | 40 | 23 | -11 | 17 | 2.61 | 0.011836 |
|  |  | VTA |  | R | 3 | -13 | -10 | 16 | 2.76 | 0.008136 |
|  |  | Insula | 13 | R | 36 | 14 | -8 | 7 | 2.68 | 0.010094 |
|  |  | Insula | 47 | R | 40 | 20 | -14 | 4 | 2.55 | 0.014035 |
|  |  | Insula | 47 | R | 36 | 26 | -8 | 4 | 2.63 | 0.011298 |
| Negative |  | Temporal cortex | 22 | R | 63 | -34 | 19 | 179 | -3.08 | 0.003413 |
|  |  | Temporal cortex | 21 | L | -63 | -36 | 1 | 53 | -3.04 | 0.00381 |
|  |  | Temporal cortex | 39 | R | 42 | -49 | 25 | 46 | -2.76 | 0.008204 |
|  |  | DMN | 10 | R | 3 | 62 | 10 | 38 | -2.91 | 0.005434 |
|  |  | ACC | 32 | R | 6 | 38 | 16 | 26 | -2.90 | 0.005549 |
|  |  | DMN | 10 | R | 6 | 65 | 22 | 23 | -2.81 | 0.007161 |
|  |  | ACC | 9 | R | 3 | 44 | 19 | 9 | -2.72 | 0.008941 |
|  |  | Temporal cortex | 39 | R | 39 | -57 | 28 | 2 | -2.52 | 0.015008 |
|  |  | ACC | 9 | R | 9 | 38 | 19 | 1 | -2.51 | 0.015254 |

The main effect of *Age* was associated with activation across occipito-temporal regions, mainly in the visual association regions (BA 18) as well as limbic regions (PHG, amygdala) and a parietal cluster (figure 2B). Random effects GLM maps of the *Age* contrasts (infancy>childhood, infancy>young adulthood, figure S3), show that visual association regions are activated across all contrasts. There was considerable resemblance between the Adulthood>Infancy and Childhood>Infancy contrast maps, while in the childhood>Adulthood contrast map the activations were weaker and sparse. Critically, there was no significant interaction between *Self-relatedness* and *Age* at the whole brain level (figure 2).

Next, we examined the preregistered regions of interest to test our hypothesis of a network responding to the Relational Self. Beta weights were extracted from ROIs and analyzed with a 2×3 (*Self-relatedness* (*Self/Other*) \**Age* (*Infancy/Childhood/Young adulthood*)) repeated measures ANOVA. we examined the hypothesis that the ROIs would show higher BOLD activity when viewing *Self* videos than when viewing similar videos of others. Indeed, results revealed that across all ROIs, *Relational Self*-videos elicited stronger BOLD activity and this difference was significant in all regions except for the DMN. Significant self-relatedness effects were found in the insula (F_(1,49)_=4.56,p=0.03, η_p_^2^=0.08, BF_exc_=0.86), PHG (F_(1,49)_=5.34,p=0.02, η_p_^2^=0.09, BF_exc_=0.971), temporal cortex (F_(1,49)_=5.18,p=0.027, η_p_ ^2^=0.096 BF_exc_=0.489), ACC (F_(1,49)_=9.46,p=0.003, η_p_^2^=0.16, BF_exc_=0.273), and amygdala (F_(1,49)_=9.87,p=0.003, η_p_ ^2^=0.16, BF_exc_=0.093). The extremely low BF in the VTA (F_(1,49)_=18.34, p<0.001, η_p_^2^=0.27, BF_exc_=0.003), and Thalamus (F =19.81,p<0.001,η_p_^2^=0.28,BF_exc_=3.42*10^−5^) suggests very strong evidence for the inclusion of the self-relatedness model, whereas in the temporal cortex and insula the evidence is moderate and in the PHG BF is relatively high, suggesting very weak evidence for self-relatedness model inclusion. As predicted, our preregistered ROIS showed ‘self-preference’ – increased activation for the *Relational Self* condition compared to ‘*other-in-relationship*’ condition (figure 3). Contrary to our hypothesis, the default mode network did not show significant differential activation for the *Relational Self*-stimuli compared to Other and BF suggested inconclusive evidence for the *Self-relatedness* model (F_(1,49)_=2.69, p=0.107, BF_exc_=2.054).

**Figure 3:**
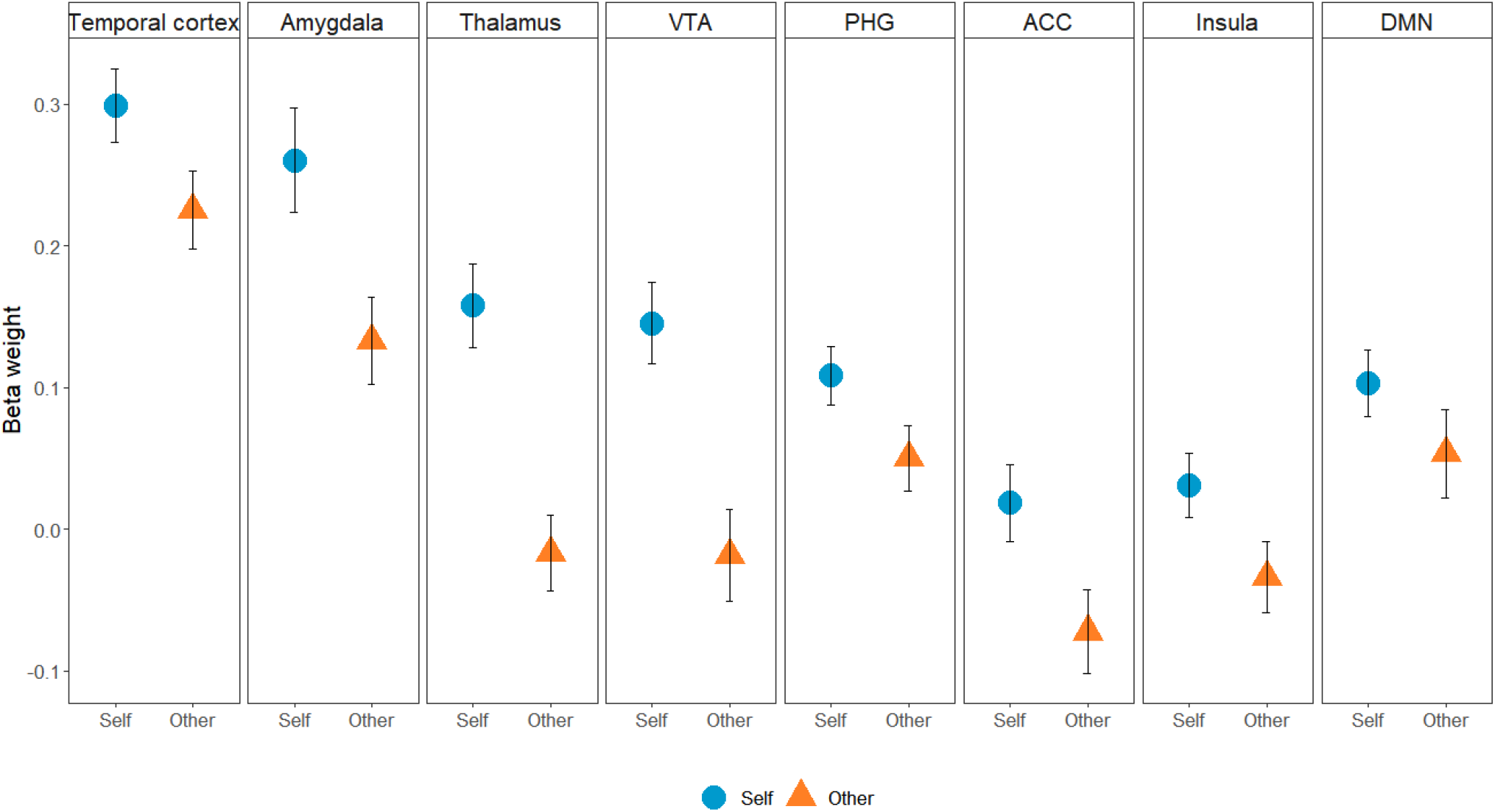
Self-relatedness main effect in ROIs, N=50. All ROIs except the DMN showed *Self* main effect. Note the strong effect in the Thalamus. Colored shapes mark the average, whiskers mark SE.

Following, we investigated if these regions would show differential responses to stimuli from the three ages (infancy/childhood/young adulthood). The temporal cortex was the only region to show a significant main effect for *Age* (F_(1.5,76)_=6.85, p=0.004, η_p_ ^2^=0.12, BF_exc_=0.003). Other regions showed strong evidence for exclusion of the age model: Insula (BF_exc_=21.71), ACC (BF_exc_=25.57), PHG (BF_exc_=15.48), Thalamus (BF_exc_=23.47), VTA (BF_exc_=18.38) and DMN(BF_exc_=12.09), in the amygdala no such strong evidence was found (BF_exc_=2.22). Thus, contrary to *Self-relatedness*, only one ROI showed sensitivity to the differences in *Age factor*.

Next, we examined interaction effect of *Self-Relatedness* X *Age conditions*. In the RM ANOVA, there was no significant interaction effect in the whole brain analysis, as well as in any of the eight preregistered ROIs (Figure S4). This finding suggests that our ROIs did not show differential Self-Other responses for different ages. However, since a lack of significant interaction effect does not provide sufficient evidence for its absence, we employed a repeated measures Bayesian ANOVA to quantify evidence for lack of such interaction (BF_exc_ factors >10 indicate strong evidence against an interaction between *Relational Self* and *Age* conditions). Bayes factors in the insula (BF_exc_=11.50), ACC (BF_exc_=81.69), PHG (BF_exc_=12.09), Amygdala (BF_exc_=11.39), Thalamus (BF_exc_=14.12), temporal cortex (BF_exc_=14.28), and DMN (BF_exc_=9.81) – all showed strong evidence for exclusion of the interaction model. In the VTA (BF_exc_=5.96), the evidence was moderate. These findings indicate that in all of our pre-registered regions (apart from the DMN), while a significant main effect for *Self-relatedness* was found, manifested as stronger neural responses to *Self* stimuli, *Age-*related effects (i.e. age differences in stimuli To capture the ongoing exchange of information between are ROIs during the task,) emerged only in the temporal cortex ROI. Critically, there was strong evidence against interaction model across all but one of our ROIs, indicating age invariant brain activity for the *Relational Self*.

To capture the ongoing exchange of information between our ROIs during the task, we utilized psycho-physiological interaction (PPI) analysis. Briefly, PPI examines task-specific changes in the correlated activity across different brain regions by identifying voxels that show increased correlation with a seed region within a given psychological context (=condition). The variance explained by the interaction term resulting from the PPI represents explained variance above and beyond the variance accounted for by the main effects of task (*Self* vs. *Other*) and physiological correlation (e.g., regions that are associated anatomically, activated by a third region etc.); Hence, significant PPI correlations represent the tightening or loosening of connectivity among regions in response to the task.

In order to investigate such fluctuations, we created a multi subject Random effect GLM of PPI from an ACC seed to our preregistered ROIs. The resulting map (cluster threshold corrected, Figure 4) indicated that the left Insular cortex had increased correlation with ACC in the *Relational Self* condition compared to the Other condition. This indicates that during the viewing of Relational Self-stimuli across the individual’s developmental history, the transfer of information between ACC and insula is stronger than in non-self stimuli.

**Figure 4:**
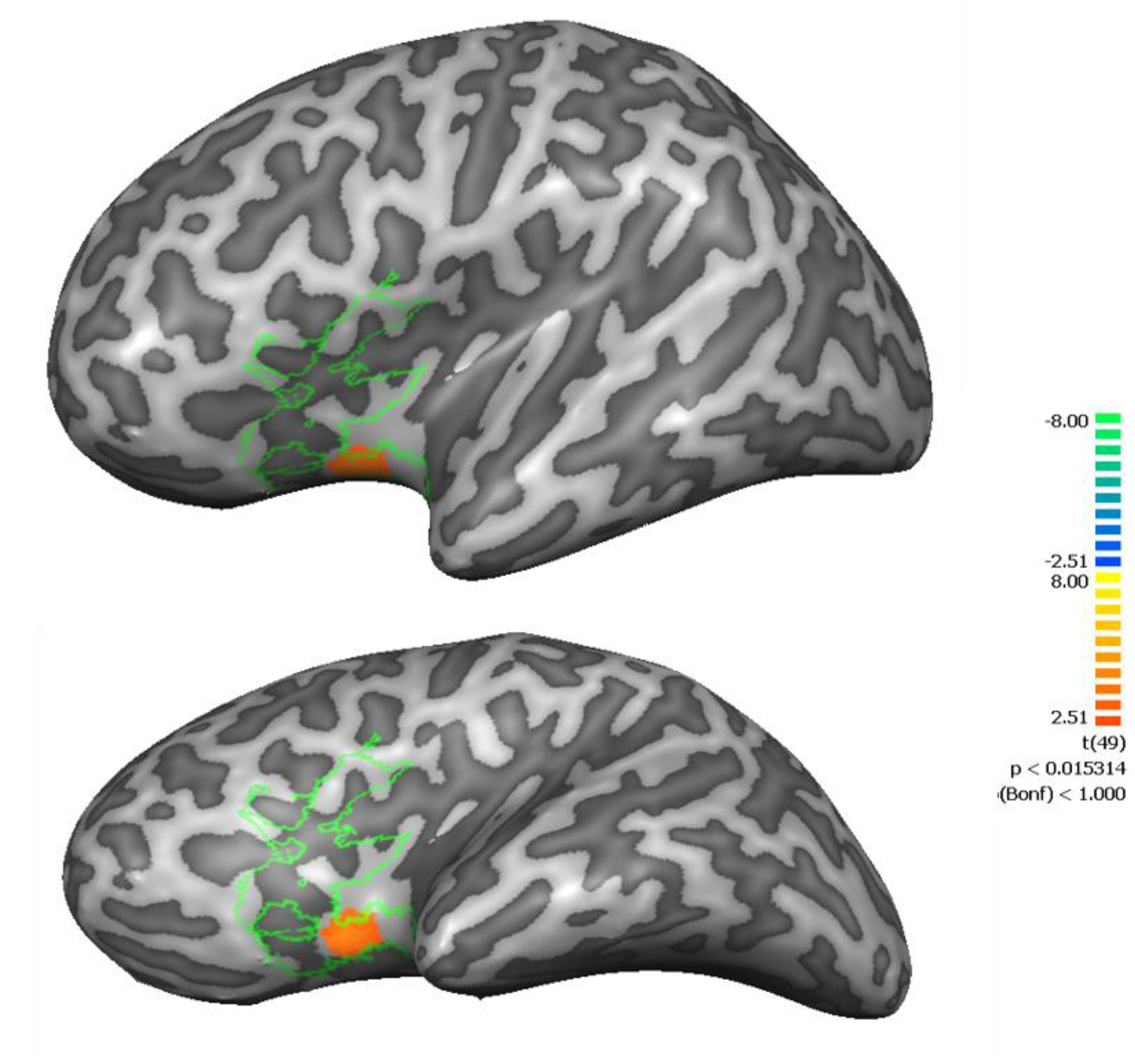
Psychophysiological interaction analysis, with ACC seed for self>other contrast of 50 participants. (analysis dataset). Map was corrected using a Monte-Carlo cluster-level estimation. A cluster in the left insula (ROI outline in green).

Finally, we explored the relationship between the magnitude of ACC and insular activation during the *Self* conditions and the independent coding of maternal and child’s social behavior during the interaction. We focused on the two main maternal and child constructs in the attachment literature: maternal sensitivity and child positive social engagement, which have repeatedly shown to be individually-stable from infancy to adolescence and to predict positive social-emotional and psychiatric outcomes. ACC beta values for *Self* conditions were positively correlated with *Child Positive Engagement* (r=0.375, p=0.007; Bonferroni corrected, figure S5), but not with *Maternal Sensitivity* (r=0.242, p=0.091). Similarly, Insula beta values for the *Self* conditions were moderately correlated with *Child Positive Engagement* (r=0.280, p=0.049; uncorrected) (figure 5), but not significantly correlated with *Maternal Sensitivity* (r=0.237, p=0.097). These findings suggest that the more the child showed initiation of social communications, positive affect, warmth, motivation, and involvement during interactions with the mother, the greater was the ACC in insula neural response to viewing *Relational Self* stimuli. Notably, no significant correlations were found between *Child Positive Engagement* and neural response to the *Other* condition in the ACC (r=0.179, p=0.168) and insula (r=0.062, p=0.667).

**Figure 5:**
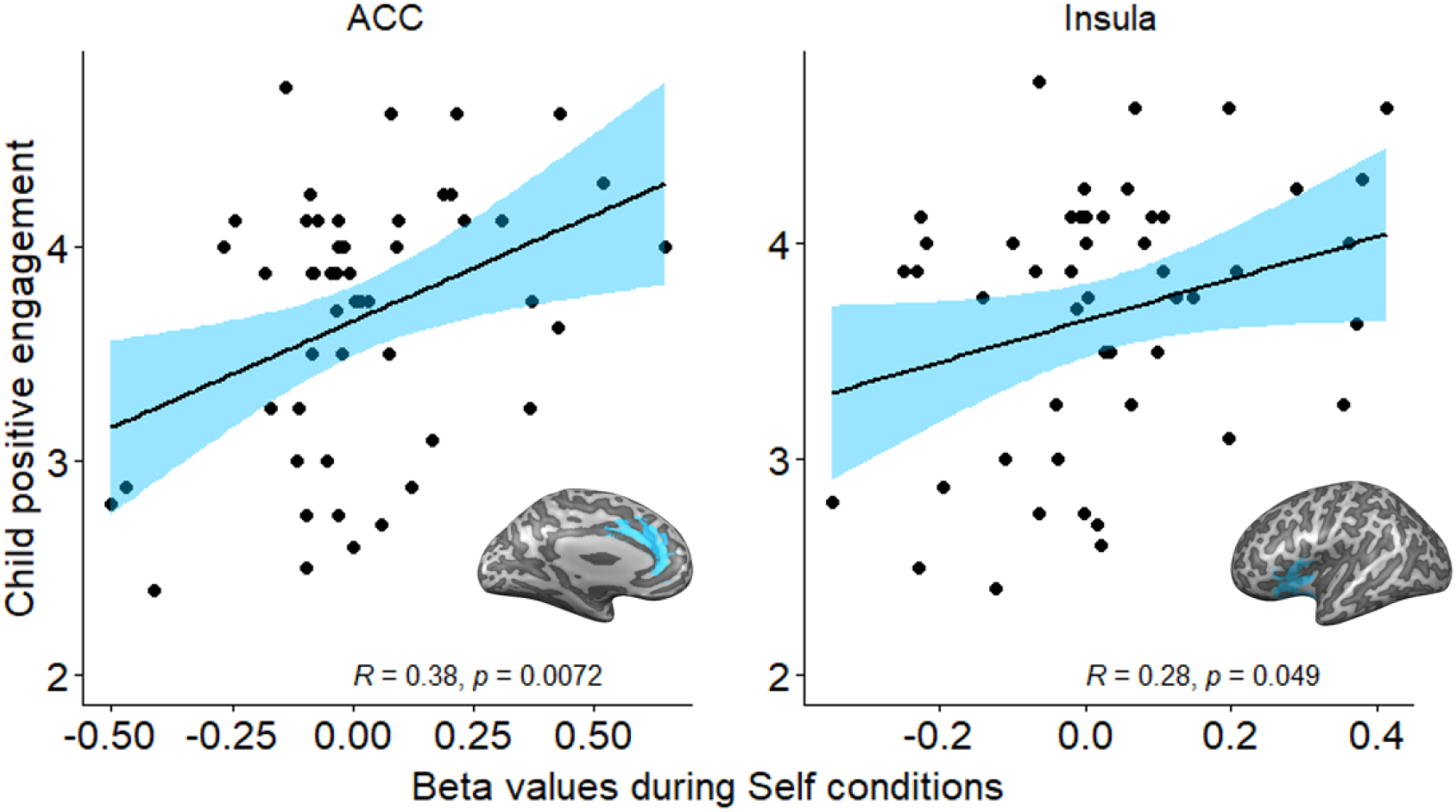
Child positive engagement correlation with activation during *Relational Self* conditions in the ACC and Insula. Pearson correlation

## Discussion

Describing the neural underpinnings of the Self poses a fundamental challenge due to the inherent complexity and multi-dimensionality of the construct and the diverse methodologies used (Blanke et al., 2015; Salomon, 2017; Frewen et al., 2020; Yeshurun et al., 2021). Our study addresses a novel aspect in research on the neural systems of the self and aims to capture its relational dimension, which develops in the context of the mother-infant attachment (Allen and Tsakiris, 2018; Ciaunica and Crucianelli, 2019; Montirosso and McGlone, 2020). In searching for the neural underpinnings of the Relational Self, our paradigm exposed young adults to video reminders of interactions with the mother across their entire relational history from infancy to adulthood. This paradigm includes several key aspects of the self previously addressed in neuroscience research (Northoff et al., 2006; Peer et al., 2015); our stimuli include aspects of visual and auditory self-recognition (Uddin et al., 2005; Qin and Northoff, 2011; Salomon et al., 2012, 2020), autobiographical memory, affective self-relevance (Kelley et al., 2002; Schäfer et al., 2020), and “self-relatedness” factors (mother’s face, mother’s voice, my home). These dimensions were encored within the “narrative self”, the “story” the individual builds of his or her personal development, attachment relationships, and childrearing environment (Gergen and Gergen, 1988; Miller et al., 1990).

Several important findings emerged from our study. First, we found that the same attachment network sustaining the parent’s attachment to the child also activates when the child observes attachment-related “Self” stimuli. While our study is among the first to assess the brain basis of attachment in the child-to-parent relationship, findings are consistent with the formulations of attachment theory (Bowlby, 1969) and research in animal models on the cross-generational transmission of the neurobiology of attachment (Meaney and Champagne, 2001). Our pre-registered ROIs showed greater activations to Relational-Self stimuli compared to similar but unfamiliar stimuli and the increased activations were in the same nodes as those found in the parents’ brain in response to their child compared to unfamiliar child (Ranote et al., 2004; Noriuchi et al., 2008; Musser et al., 2012; Feldman, 2015b; Rigo et al., 2019). These included subcortical regions implicated in mammalian caregiving, such as amygdala and hippocampus, and cortical areas, particularly ACC, insula, and temporal regions.

Second, we found that activations to the Relational Self across a twenty-year span were time-invariant and these regions showed consistently larger activations for the self across ages. Our Bayesian analysis indicated strong evidence for lack of interaction between the *Age* and *Self-relatedness* factors in the ROIs, indicating that the greater activity for Relational Self-stimuli did not differ as a function of age. This is particularly striking as the stimuli differed markedly across ages. While a three-month-old infant and a twentysomething adult interacting with the mother differ on any possible sensory and mental self-related parameter; looks (body, face), verbal content, temporal distance, self-similarity, affective expression, or mentalization, it appears that the identification of the stimuli with the “self” trumps these differences and activations in the Relational Self nodes were mainly consistent from infancy to adulthood.

Third, Relational-Self stimuli elicited not only greater activations but also increased inter-region coherence. The PPI analysis indicated that our seed region in the ventral ACC showed increased connectivity with the insula during observation of Relational Self-stimuli. Finally, the magnitude of ACC and insular activations were linked with the degree of positive engagement, motivation for social connection, and initiation of social communication the child’s exhibited during interaction with the mother across the twenty-year span from infancy to adulthood.

The ACC is an integrative interface of sensation, cognition, emotion, arousal, and neuromodulation (Peterson et al., 1999; Bush et al., 2000; Rolls, 2019; Vassena et al., 2020), is among the most interconnected hubs in the brain (Margulies et al., 2007), and is a core region sustaining self-related processes (Northoff and Bermpohl, 2004; Qin and Northoff, 2011). The ACC contains evolutionary-conserved projections to all primary sensory and associative cortices to receive exteroceptive input; reciprocal projections to amygdala, hippocampus, ventral striatum, and VTA to enable regulation of emotion and motivation; and with OFC for processing reward and prioritizing action based on valuation (Burgos-Robles et al., 2019). Extensive reciprocal projections connect the ACC with the insula into a global paralimbic interface that combines interoceptive signals from the body with exteroceptive cues into an integrative percept that enables embodiment and emotional mirroring (Craig, 2009; Seth, 2013; Park et al., 2018), which are critical for the formation of attachment bonds. Both the ACC and insula contain layer V von Economo neurons that afford rapid communication among the two regions as well as with other upstream or downstream targets (Allman et al., 2011). Both regions contain areas of overlap between self and others’ pain (Corradi-Dell’Acqua et al., 2016; Smith et al., 2021), which mark them as regions for the interplay of connection and separation of self and other that define the basis of close relationships throughout life.

It has long been known (Papez, 1937) that the ACC plays a key role in affective processing which underpins the formation of attachment bonds. The ACC is implicated in evaluative emotional processing (Esslen et al., 2004), assessment of the motivational value of stimuli (Fujiwara et al., 2009), generation of behavioral emotional response (Etkin et al., 2011), and top-down monitoring of affective information (Carter et al., 2001). Furthermore, the ACC integrates social functions relevant for the Relational Self. It contains neurons that respond specifically to information related to self versus non-self (Sturm et al., 2013) and integrates sensory, cognitive, and affective information into a coherent percept that prioritizes motivation (Porter et al., 2019; Lee and Reeve, 2020), forms predictions, regulates affect (Ochsner et al., 2009), consolidates memories (Restivo et al., 2009; Vetere et al., 2011), and shapes the individual’s mode of operation in social contexts (Krill and Platek, 2009; Vassena et al., 2017). Such integration of functions related to self, affect, and social processing renders the ACC a key region for the Relational Self. Our results on the increased connectivity between the ACC and insula suggest that this interface becomes more functionally coupled in response to own attachment stimuli to chart the neural underpinnings for the Relational Self.

Several meta-analyses (Northoff et al., 2006; Qin et al., 2020) have shown that midline cortical structures, including our seed region in the ventral ACC, provide a functional network that integrates regions across the ventral and dorsal midline to sustain the multiple dimensions of the self, including proto-self, self-”qualia”, bodily self, facial-self recognition, and mental self (Northoff and Bermpohl, 2004; Uddin et al., 2007; Northoff and Panksepp, 2008; Moran et al., 2009). The ventral part of the midline cortical structures includes an area of overlap among the post-genual ACC, our seed region, the vmPFC, and the mOFC, and is particularly linked with self-referentiality and the narrative self (Northoff et al., 2006; Araujo et al., 2013; Salomon et al., 2014). A recent meta-analysis (Qin et al., 2020) differentiated on the basis of all available brain studies of the Self between three levels; the bodily, which involves the interoceptive monitoring of one’s body, the environmental, that includes self-relevant exteroceptive signals, and the mental, which represents non-material self-components. While the insula is represented at all three levels of the Self, the ventral ACC is implicated only in the mental level. This level expands the representations of the Self beyond the bodily or immediate sensory into autobiographical memory, personal perspective, and self-reference.

The coupling between the ACC and insula during observation of Relational Self-stimuli anchors the representation of the mother-child attachment in the bodily and early non-verbal sensory but integrates these interoceptive levels into an adult representation (Morita et al., 2014). Such mental Relational Self can reverberate the entire primary relationship and prepares the child for other attachments throughout life, with romantic partners, friends and eventually with the child’s own children. Insular activations in the parental brain were found to correlate with behavioral markers of the mother-child relationship (Abraham et al., 2016) and maternal insular activation are thought to provide an external regulation of the infant’s emerging capacities recognize his/her own bodily signals and develop interoceptive representations (Atzil et al., 2018). As part of the sociotemporal brain (Schirmer et al., 2016), the insula also monitors the duration and patterns of the early mother-infant synchronous interactions, which later expand in the dyadic relationship and are individually stable from infancy to adulthood (Ulmer-Yaniv et al., 2020). Such cingulate-insula connectivity provides the basis for maternal caregiving of the infant (Kim et al., 2010; Swain et al., 2017; Rigo et al., 2019) and, as seen by our PPI, also underpins the neural representation of the time-invariant Relational Self.

Interoception, the neural representation of one’s own internal bodily signals, is an important feature of the Relational Self, particularly with regards to the mother-child attachment. Recent models on interoception (Chen et al., 2021) suggest that viewing the self often triggers activation in nodes of the interoceptive network, and own mother-infant interaction may be a particularly strong reminder of caregiving and bodily contact. Interoceptive information is first processed in the brainstem nucleus of the solitary tract and then projects to the thalamus, from where it is relayed to higher targets; the amygdala, insula, and ACC. The insula is a key hub of interoception and its rapid reciprocal projections with ACC integrates interoceptive signals with exteroceptive information into higher-order representation (Craig, 2003; Salomon et al., 2016). Recent work has highlighted links between insular interoceptive-related activity and self-related processing at both the bodily (Salomon et al., 2016; Park et al., 2018; Park and Blanke, 2019) and narrative levels (Babo-Rebelo et al., 2016; Tallon-Baudry et al., 2018). As seen (Figure 2), nodes of the interoceptive network were found here to differentiate Self versus Other; thalamus-to-brainstem, amygdala, hippocampus, insula, and ACC. This suggests some overlap between the Relational Self and the identification of signals from one’s body, indicating that the development of the sense of self and its bodily dimension develops within the context of the mother-infant bond.

Several study limitations should be considered. First, similar to all studies of the Self, it is possible that self-related stimuli are allotted more attentional resources than non-self-relevant stimuli. However, this attentional account would also suggest that the novel Self stimuli from the earlier ages would probably elicit greater attention; still our data show no difference between ages, suggesting that these findings do not stem from differential attentional engagement. Second, stimuli presentation-order was counterbalanced for *Self-relatedness*, but not for *Age* presentation order. This stemmed from our desire to present a coherent narrative account of the mother-child relationship from infancy to adulthood and describe the unfolding of the Relational Self. Still, the lack of counterbalance in age is a study limitation. Additionally, as in all ecological studies, our stimuli varied on numerous visual and auditory properties. Despite these limitations, the Self-related effects found in our preregistered ROI speak to the robustness of the effect above and beyond the idiosyncrasy of specific stimuli. Much further research is needed to characterize the development of the self-within-attachment relationships and understand the impact of culture, context, habit, and risk conditions on maturation of the Relational Self across time and within the individual’s attachment bonds throughout life.

## Acknowledgements

We would like to thank the mothers and children for their participation and cooperation.

## Supplementary materials

**Figure S1:**
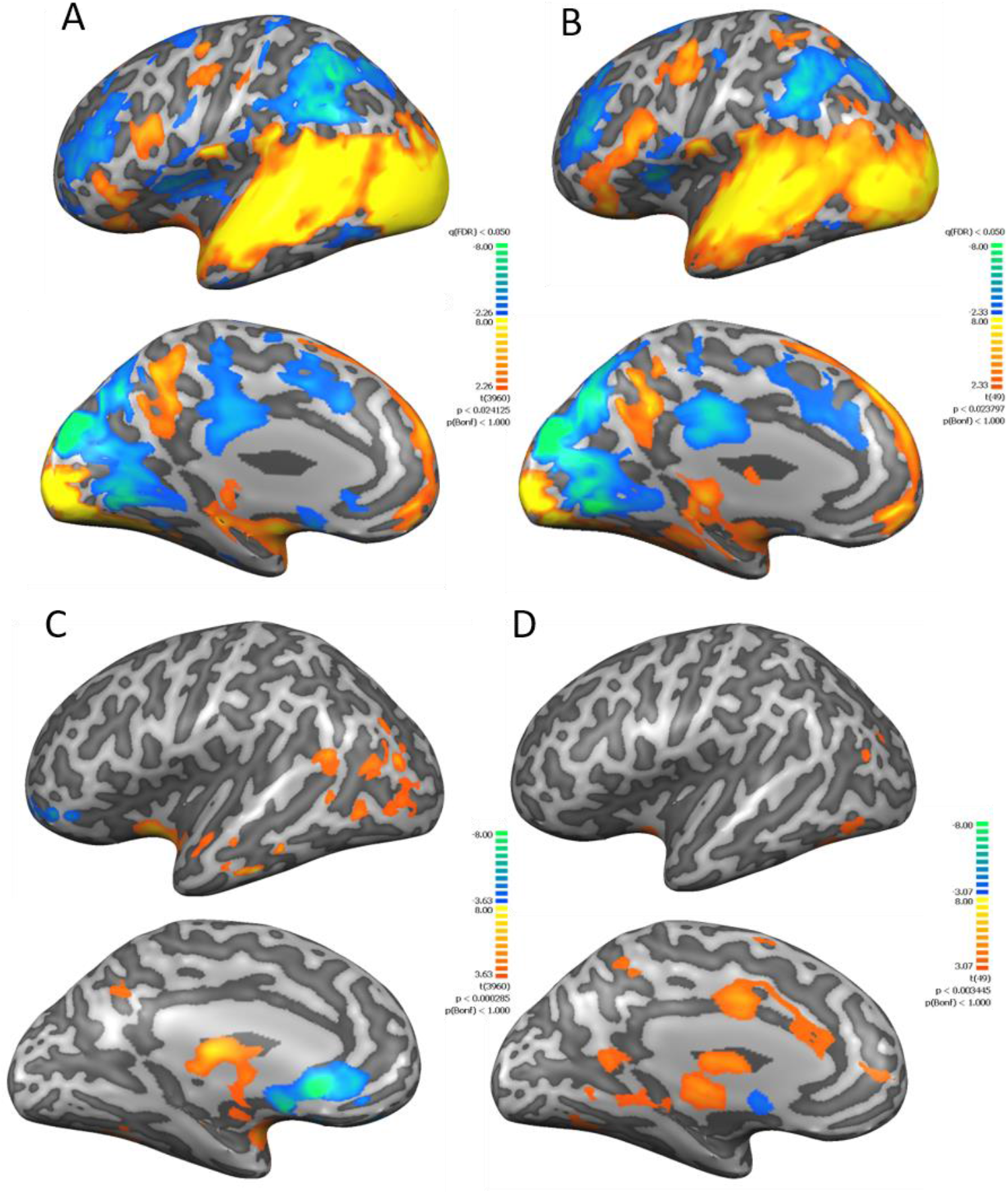
Whole brain multi subject GLM maps. **A**: Cohort 1 dataset (N=15), fixed effects GLM, whole brain all visual map, corrected for multiple comparisons at q(FDR)<0.05 **B**: Analysis dataset (N=50), random effects GLM, whole brain all visual map, corrected for multiple comparisons at q(FDR)<0.05.**C**: Cohort 1 dataset (N=15), fixed effects GLM, whole brain self>other map corrected for multiple comparisons below q(FDR)<0.05.**D**: Analysis dataset (N=50), random effects GLM, whole brain self>other map, uncorrected.

**Figure S2:**
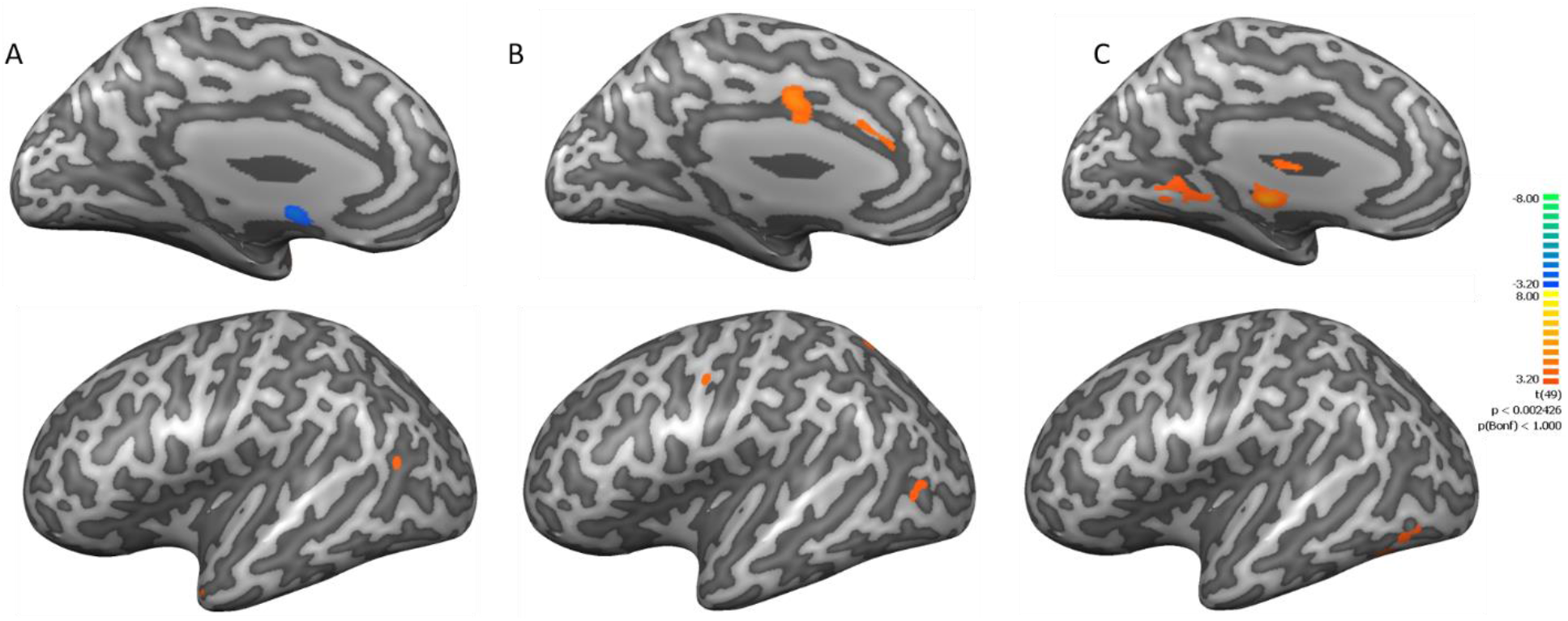
Whole brain multi subject random effects GLM maps, analysis dataset (N=50) self>other contrast for each timepoint: **A**: Infancy **B**: Childhood **C**: Adulthood. Maps are uncorrected.

**Figure S3:**
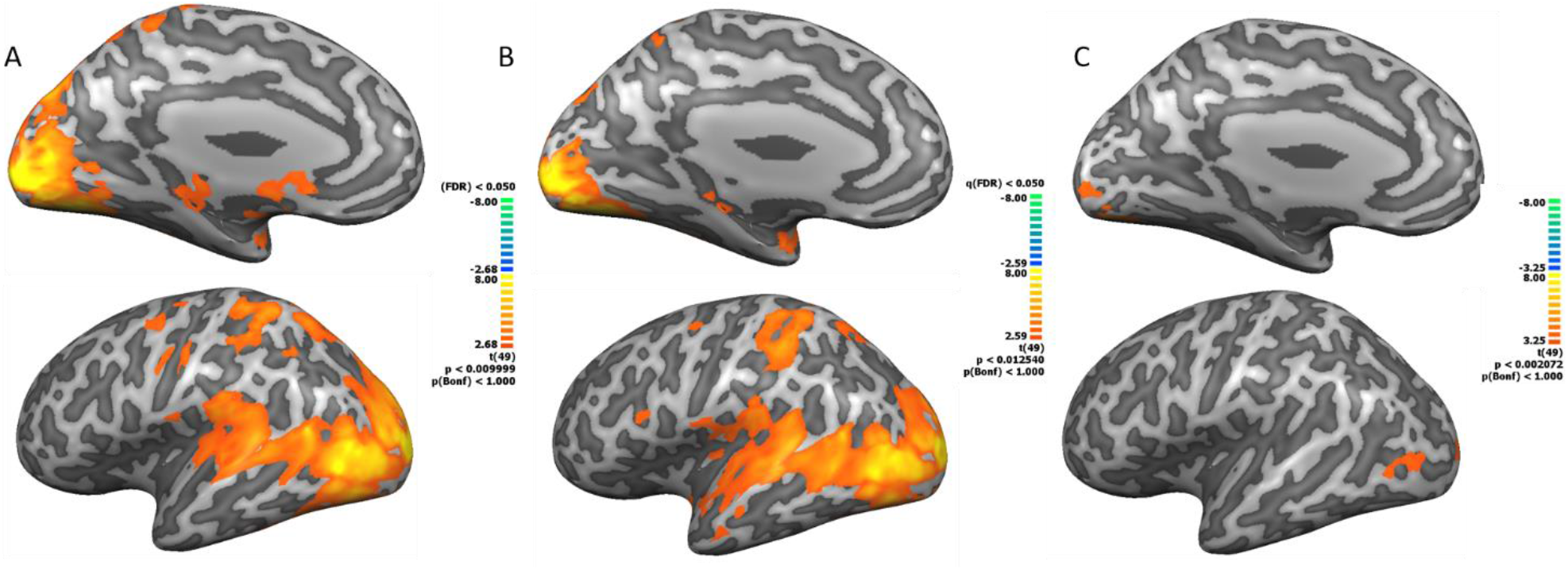
Time contrasts. whole brain multi subject random effects GLM maps, analysis dataset (N=50). **A. Infancy>childhood**, corrected for multiple comparisons at q(FDR)<0.05 **B. Infancy>young Adulthood**, corrected for multiple comparisons at q(FDR)<0.05 **C. Childhood>Young adulthood**, uncorrected.

**Figure S4:**
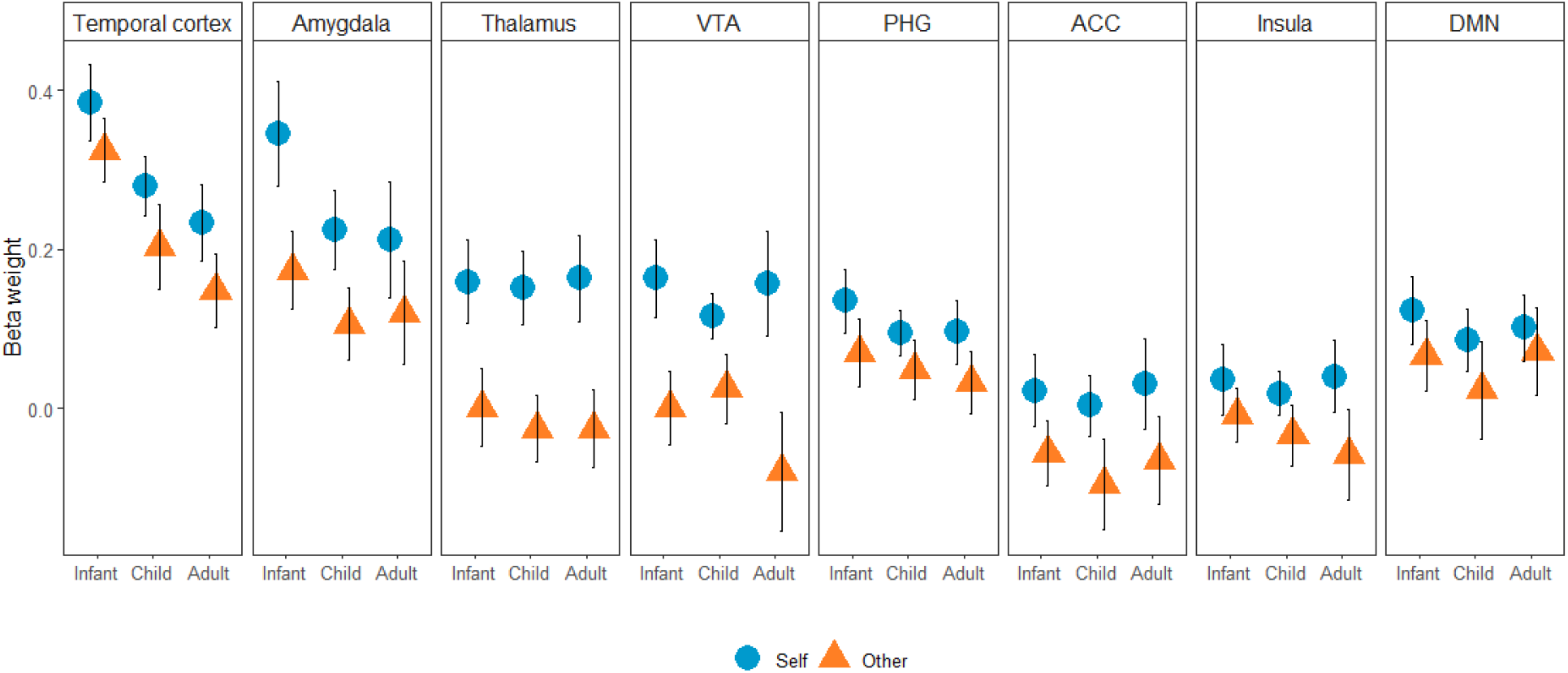
*Self-relatedness* and *Age* effects in ROIs. All ROIs except the DMN showed self-relatedness main effect. Note the strong *Self relatedness* effect in the Thalamus and *Age* main effect in the Temporal cortex. There was no interaction effect in any of our ROIs. Colored shapes mark the average, whiskers mark SE.

